# Exploring protein orthogonality in immune space: a case study with AAV and Cas9 orthologs

**DOI:** 10.1101/245985

**Authors:** Ana M. Moreno, Nathan Palmer, Fernando Alemán, Genghao Chen, Andrew Pla, Wei Leong Chew, Mansun Law, Prashant Mali

## Abstract

A major hurdle in protein-based therapeutics is the interaction with the adaptive immune system, which can lead to neutralization by circulating antibodies and clearance of treated cells by cytotoxic T-lymphocytes. One method of circumventing these issues is to use human or humanized proteins which avoid the immune response by self-recognition. However, this approach limits potential protein therapeutics to those of human origin, excluding many exciting effectors and delivery vehicles such as CRISPR-Cas9 and adeno-associated viruses (AAVs). To address this issue, we propose here the sequential use of orthologous proteins whose function is constrained by natural selection, but whose structure is subject to diversification by genetic drift. This would, in principle, allow for repeated treatments by ‘immune orthogonal’ orthologs without reduced efficacy due to lack of immune cross-reactivity among the proteins. To explore and validate this concept we chose 91 Type II CRISPR-Cas9 orthologs and 167 AAV capsid protein orthologs, and developed a pipeline to compare total sequence similarity as well as predicted binding to class I and class II Major Histocompatibility Complex (MHC) proteins. Interestingly, MHC binding predictions revealed wide diversity among the set of Cas9 orthologs, with 83% of pairs predicted to have non cross-reacting immune responses, while no global immune orthogonality among AAV serotypes was observed. To confirm these findings we selected two Cas9 orthologs, from *S. pyogenes* and *S. aureus*, predicted to be orthogonal in immune space, and delivered them into mice via multiple AAV serotypes. We observed cross-reacting antibodies against AAV but not Cas9 orthologs in sera from immunized mice, validating the computationally predicted immune orthogonality among these proteins. Moving forward, we anticipate this framework can be applied to prescribe sequential regimens of immune orthogonal protein therapeutics to circumvent pre-existing or induced immunity, and eventually, to rationally engineer immune orthogonality among protein orthologs.

## INTRODUCTION

Protein therapeutics, including protein-based gene therapy, have several advantages over small-molecule drugs. They generally serve complex, specific functions, and have minimal off-target interference with normal biological processes. However, one of the fundamental challenges to any protein-based therapeutic is the interaction with the adaptive immune system. Neutralization by circulating antibodies through B-cell activation and clearance of treated cells by CD8+ cytotoxic T-lymphocytes (CTLs) create a substantial barrier to effective protein therapies^1–4^. Although the delay in the adaptive immune response to novel proteins may allow sufficient time for the initial dose to work, subsequent doses face faster and stronger secondary immune responses due to the presence of memory T-and B-cells. In addition, gene transfer studies have shown that host immune responses against the delivery vector and/or therapeutic transgene can eliminate treated cells, thus limiting the efficacy of the treatment^5–10^.

A common approach to circumventing these issues has been to utilize human proteins, or to humanize proteins by substitution of non-human components^11,12^. However, this approach is limited to a small set of therapeutic proteins naturally occurring in humans or closely related species. In addition, although the humanization of proteins can result in a significantly less immunogenic product, they still carry immunological risk^12^. Another way to circumvent an immune response to protein therapeutics is the removal of immunogenic T cell epitopes^13,14^. Once immunogenic T cell epitopes are identified, substitution of key amino acids may reduce the protein’s immunogenicity since modification of amino acids at critical anchor residues can abrogate binding to MHC molecules and prevent antigen presentation. However, this can prove difficult due to the massive diversity at HLA loci. As epitope engineering must account for the substrate specificity of each different HLA allele, therapeutics would likely have to be uniquely modified for each patient. All the same, epitope deletion has been successfully applied to several proteins^15^, but can only preserve protein function when limited to small numbers of HLA alleles unrepresentative of the full diversity. Structural modifications such as PEGylation have also been known to reduce immunogenicity by interfering with antigen-processing mechanisms. However, there is evidence that PEG-specific antibodies are elicited in patients treated with PEGylated therapeutic enzymes^16–19^.

Furthermore, protein therapies have required repeated treatments due to degradation of the protein or turnover of treated cells, or, in the case of gene therapy, reduced expression of the transgene^20,21^. This provides an even greater challenge as repeated exposure to the same antigen can elicit a more robust secondary immune response^22^, which may completely inhibit subsequent dosage or even sensitize the immune system to antigens remaining from the initial exposure. In order to facilitate efficacious repeat protein therapies, we propose the use of orthologous proteins whose function is constrained by natural selection, but whose structure is subject to diversification by genetic drift. An ortholog, given sufficient sequence divergence, will not cross-react with the immune response generated by exposure to the others, allowing repeat doses to avoid neutralization by existing antibodies and treated cells to avoid clearance by activated CTLs.

As a case study for exploring this approach we focused on the CRISPR-Cas9 system, perhaps the most anticipated therapeutic for gene editing^23–33^. Comparative genomics has demonstrated that Cas9 proteins are widely distributed across bacterial species and have diversified over an extensive evolutionary history^34–37^. Although there may be pre-existing immunity to Cas9s from some commensal species^38^, we hypothesized this diversity could provide a mechanism to circumvent inducing immunological memory by utilizing orthologous Cas9 proteins for each treatment. Additionally, the immunogenicity due to the delivery vehicle or administration route for the Cas9 and the associated guide RNA (gRNA) must also be considered. In this regard, adeno-associated viruses (AAVs) have emerged as a highly preferred vehicle for gene delivery, as these are associated with low immunogenicity and toxicity^8,9^, which promotes long-term transgene expression^39,40^ and treatment efficacy. Despite the relatively low immunogenicity of AAV vectors, antibodies against both the capsid and transgene may still be elicited^41–46^. Additionally, the prevalence of neutralizing antibodies (NAB) against AAVs in the human population^47^ and cross-reactivity between serotypes^48^ remains a hurdle for efficacious AAV therapy. Although AAVs were initially considered non-immunogenic due to their poor transduction of antigen-presenting cells (APCs)^49^, it is now known that they can transduce dendritic cells (DCs)^50^ and trigger innate immune responses through Toll-like receptor (TLR) signaling pathways^51^. The ability to transduce DCs is dependent on AAV serotype and genome, and may be predictive of overall immunogenicity^52^.

To evaluate the immune orthogonality of AAV-delivered CRISPR-Cas systems, we analyzed 91 Cas9 orthologs, and 167 AAV VP1 orthologs. By comparing total sequence similarity as well as predicted binding strengths to class I and class II MHC molecules, we constructed graphs of immune cross-reactivity and computed cliques of proteins that are orthogonal in immunogenicity profiles. Although MHC epitopes do not predict antibody epitopes, the induction of the more powerful memory response is primarily dependent on reactivation of memory B-cells with help from memory T-cells through the presentation of antigens on class II MHC molecules^53,54^. Finally, we experimentally confirmed our immunological predictions by assaying treated mice for induction of protein-targeting antibodies.

## RESULTS

### Humoral immune response to AAV and Cas9

One of the major obstacles for sequential gene therapy treatments is the presence of neutralizing antibodies against the delivery vehicle and transgene cargo induced by the first administration of the therapy. To determine the humoral immune response kinetics to the AAV-8 capsid and the Cas9 transgene, we first injected C57BL/6J mice retro-orbitally with 10^12^ vg of AAV-8-SaCas9 targeting proprotein convertase subtilisin/kexin type 9 (PCSK9), a promising gene target that when disrupted can reduce Low Density Lipoprotein (LDL) levels and protect against cardiovascular disease. Consistent with a previous study^55^, mice had reduced PCSK9 serum levels as early as one week post-injection due to successful SaCas9 mediated gene-editing, which was sustained for the entire duration of the experiment (4 weeks) **(Figure 1C)**. We noted that a subset of the mice developed IgG1 antibodies against the SaCas9 protein **(Figure 1D)**. Additionally, mice developed humoral immunity to the AAV8 capsid withing one week post-injection **(Figure 1E)**. To evaluate the feasibility of multiple dosing with AAV-Cas9, we next investigated whether immune orthogonal sets of AAV and Cas9 orthologs exist.

**Figure 1.**
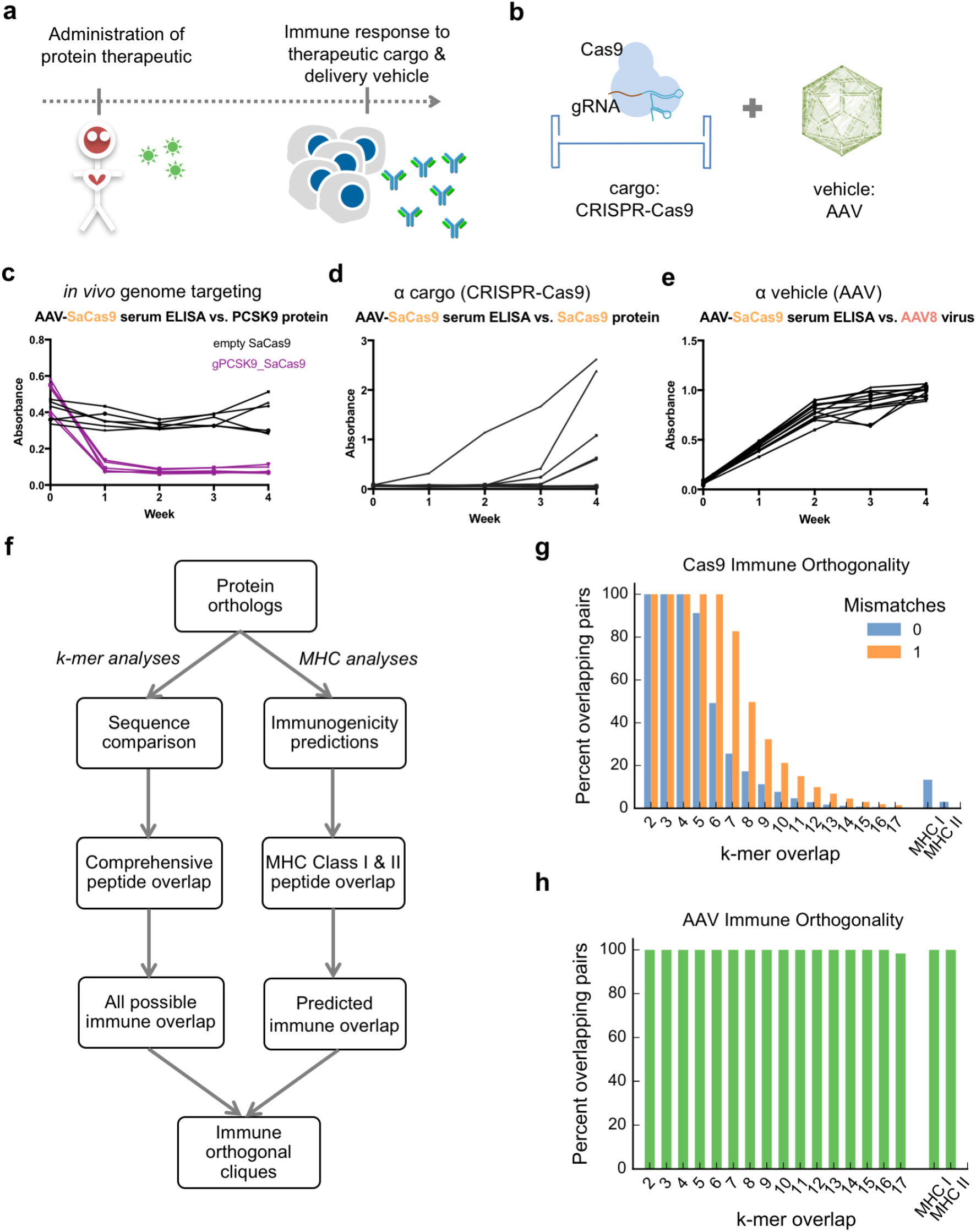
Protein based therapeutics elicit an adaptive immune response: experimental and *in silico* analyses. **(a)** Proteins have substantial therapeutic potential, but a major drawback is the immune response to both the therapeutic protein and its delivery vehicle. **(b)** As a case study, we explored the CRISPR-Cas9 systems and corresponding delivery vehicles based on AAVs. **(c)** Mice were injected retro-orbitally with 10^12^ vg/mouse of AAV8-SaCas9 targeting the PCSK9 gene or a non-targeting control (empty vector). A decrease in PCSK9 serum levels, due to successful gene targeting, can be seen in mice receiving AAV-SaCas9-PCSK9 virus (n=6 mice for each group). **(d)** Immune response to the payload was detected in ELISAs for the SaCas9 protein. (n=12) **(e)** Immune response to the delivery vehicle was detected in ELISAs for the AAV8 virus capsid (n=12 mice). **(f)** *In silico* workflow used to find immune orthogonal protein homolog cliques. **(g)** Immunologically uninformed sequence comparison was carried out by checking all *k*-mers in a protein for their presence in another protein sequence with either zero or one mismatch. The x-axis corresponds to *k*, while MHC I and MHC II show overlap only of peptides predicted to bind to MHC class I and class II molecules. 48% of Cas9 pairs show no 6-mer overlap, and 83% of pairs show no overlapping MHC-binding peptides. **(h)** Same as (g) but for AAV VP1 capsid proteins. All AAV pairs contain overlapping MHC-binding peptides.

### Identifying immune-orthogonal proteins

Natural selection produces diverse structural variants with conserved function in the form of orthologous genes. We assayed the relevance of this diversity for immunological cross-reactivity of 91 Type II Cas9 orthologs and 167 AAV orthologs by first comparing their overall amino acid sequence similarities, and second, using a more specific constraint of how their respective amino acid sequences are predicted to bind MHC Type I and II molecules **(Figure 1F)**. From these analyses we obtained first an estimate of the comprehensive immune overlap among Cas9 and AAV orthologs based purely at the sequence level, and second a more stringent estimate of predicted immune overlap based on predicted MHC binding. By sequence-level clustering and clique finding methods, we defined many sets of Cas9 orthologs containing up to 9 members with no 6-mer overlap **(Figure S1)**. Notably, based on MHC-binding predictions, we find among the set of Cas9 orthologs that 83% of pairs are predicted to have non cross-reacting immune responses, i.e. they are predicted to be orthogonal in immune space **(Figure 1G)**. On the contrary, among AAV capsid (VP1 protein) orthologs we did not find full orthogonality up to the 14-mer level, even when restricting predictions with MHC-binding strengths **(Figure 1H)**, likely reflecting the strong sequence conservation and shorter evolutionary history of AAVs^56^. This analysis suggests, consistent with previous observations^57,58^, that exposure to one AAV serotype can induce broad immunity to all AAVs, which presents a significant challenge to AAV delivery platforms, as some serotypes are prevalent in human populations. Despite the most divergent AAV serotype (AAV-5) showing the fewest shared immunogenic peptides, there remain tracts of sequences fully conserved within the VP1 orthologs. As expected, predicted immune cross-reaction negatively correlates with phylogenetic distance **(Figure S2)**, though there is significant variation not captured by that regression, suggesting that MHC-binding predictions can refine the choice of sequential orthologs beyond phylogenetic distance alone.

### Confirming humoral immune-orthogonality among Cas9 proteins

To test our immunological predictions and to establish the utility of this approach, we narrowed in on a 5-member clique containing the ubiquitously used *S. pyogenes* Cas9 in addition to the well-characterized *S. aureus* Cas9 **(Figure S1)**. To determine whether either of these proteins have cross-reacting antibody responses, we injected mice with 10^12^ vg of either AAV8 or AAVDJ capsids containing SaCas9 or SpCas9 transgenes via retro-orbital injections and harvested serum at days 0 (pre-injection), and periodically over 4–6 weeks **(Figure 2A)**. SpCas9-specific antibodies were detected in the plasma of all mice injected with SpCas9 (n=6), and notably none of the mice injected with SaCas9 (n=12) **(Figure 2B)**. Although SaCas9 appeared to induce a weaker response, as only half of the mice injected with SaCas9 AAVs (n=12) developed detectable antibodies against SaCas9, none of the mice injected with SpCas9 AAVs (n=6) developed an antibody response against SaCas9. These results were confirmed in an independent study in which SpCas9-specific antibodies, but not SaCas9-specific antibodies, were detected in the plasma of mice injected with AAV-SpCas9 (n=12). These mice were injected retro-orbitally with 10^12^ vg of AAV8-SpCas9 or AAVDJ-SpCas9, and also received an additional intramuscular injection with 10^1^1 vg at week 4. **(Figure 2C)**. Taken together, our data confirms that SpCas9 and SaCas9 have humoral immune-orthogonality.

**Figure 2.**
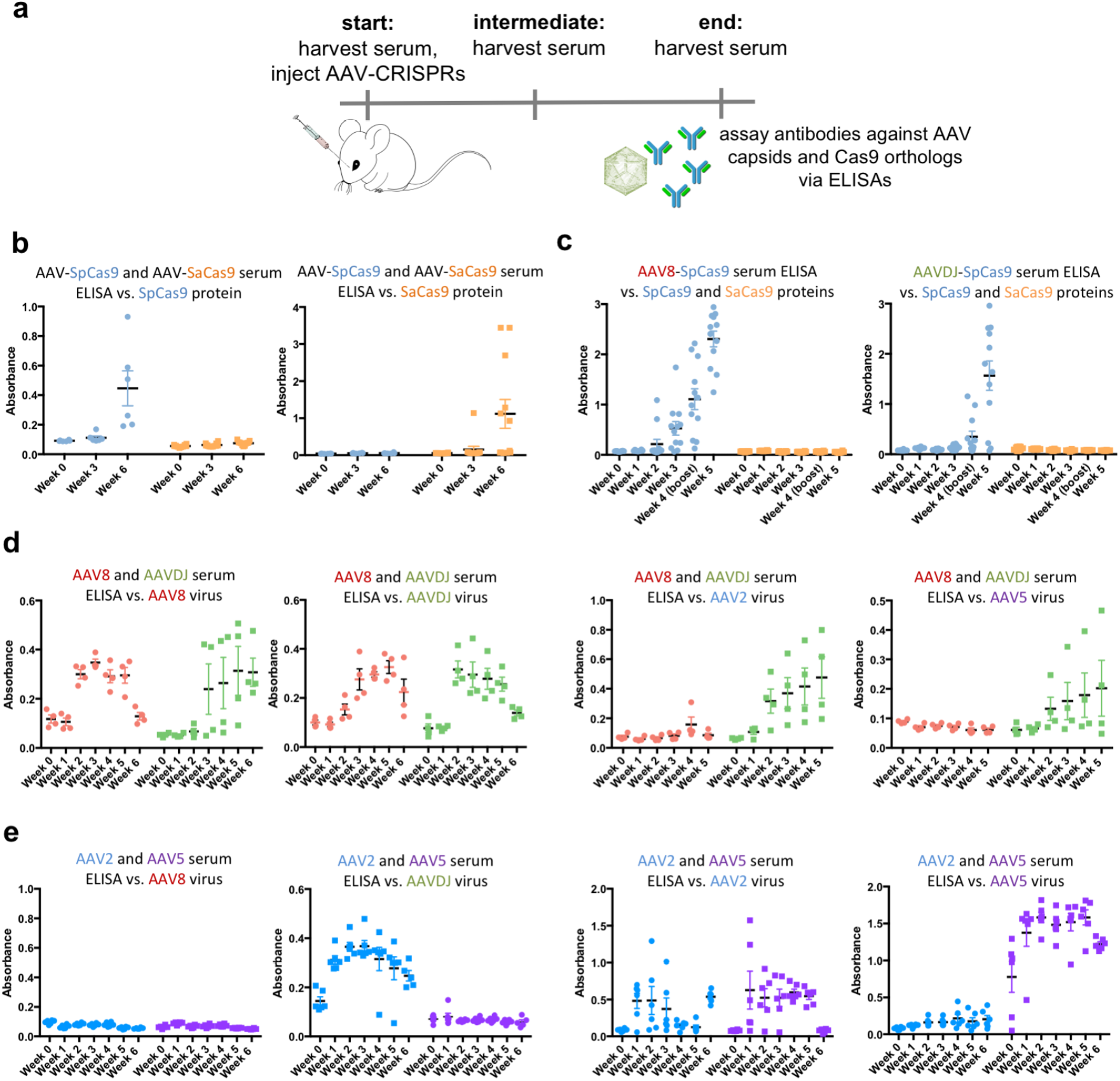
Experimental validation of Cas9 and AAV immunogenicity predictions. **(a)** Mice were exposed to antigens via retro-orbital injections at 10^12^ vg/mouse. Serum was harvested prior to injection on day 0, and at multiple points over the course of 4–6 weeks. **(b)** anti-SpCas9 antibodies generated in mice injected with SpCas9 (n=6) and SaCas9 (n=12), and anti-SaCas9 antibodies generated in mice injected with SpCas9 (n=6) and SaCas9 (n=12). **(c)** anti-SpCas9 and anti-SaCas9 antibodies generated by mice injected with AAV8 SpCas9 (n=12; left panel), or AAVDJ SpCas9 (n=12; right panel). **(d)** anti-AAV8/DJ/2/5 antibodies generated against mice injected with AAV8 or AAVDJ (n=4 for all panels). **(e)** anti-AAV8/DJ/2/5 antibodies generated against mice injected with AAV2 or AAV5 (n=5 for all panels).

### Broad cross-reactivity among AAV serotypes

AAVs are becoming a preferred delivery vehicle due to their ability to avoid induction of a strong CD8+ T-cell response, however, the presence of neutralizing antibodies remains a significant barrier to successful application of AAV therapies. Consistent with previous results^57^, we found shared immunogenic peptides among all the various human AAV serotypes, **(Figure S3)**. We confirmed the lack of orthogonality for two serotypes, AAV8 and AAVDJ, in which we found that antibodies produced in mice injected with AAV8 and AAVDJ react to both AAV8 and AAVDJ antigens **(Figure 2D)**. Our analysis suggests that there are no two known AAVs for which exposure to one would guarantee immune naïveté to another across all HLA genotypes. However, immune cross-reaction could be minimized through the use of AAV5^58,59^, the most phylogenetically divergent serotype. Our predictions identify only a single shared highly immunogenic peptide between AAV5 and the commonly used AAV2 and AAV8 in the mouse model (though several other shared peptides of mild MHC affinity exist). We confirmed this via ELISAs, where mice injected with AAV2 did not elicit antibodies against AAV5 and AAV8, and mice injected with AAV5 did not elicit antibodies against AAVDJ and AAV8 **(Figure 2E)**.

## DISCUSSION

The use of protein therapeutics requires ways to evade the host’s immune response. Cas9, as an example, has prokaryotic origins and can evoke a T-cell response, which may lead to clearance of transduced cells. In addition, circulating antibodies can neutralize the AAV vector and prevent efficient transduction upon repeated doses. Immunosuppressive drugs could mitigate some of these aspects, but not without significant side-effects, as well as not being applicable to patients in poor health^60–63^. Similar to what has been done in cancer antibody therapeutics^64^, the SpCas9 protein could also be de-immunized by swapping high-immunogenicity domains. This is a promising approach, however, it will be complex and laborious as we anticipate tens of mutations to achieve stealth, and could result in a reduction in activity and an overall less effective therapy.

To circumvent this issue, we developed here a framework to compare protein orthologs and their predicted binding to MHC I and MHC II by checking a sliding window of all k-mers in a protein for their presence in another, focusing on peptides predicted to bind to at least one MHC allele. Through this analysis, we identified cliques of Cas9 proteins that are immune orthogonal. Based on these predictions, specific T-cell responses from one ortholog would not cross-react with another ortholog of the same clique, preventing the re-activation of CD8+ cytotoxic T-cells, as well as the CD4+ T-cell help necessary to re-activate memory B-cells. We confirmed these results through ELISAs, and verified two well-characterized Cas9 proteins to be immune orthogonal, SpCas9 and SaCas9. Therefore, we expect that proteins belonging to the same clique can be used sequentially without eliciting memory T-and B-cell responses.

Due to the importance of AAVs as a delivery agent in gene therapy, we also analyzed AAV serotypes through our MHC I and II comparison framework, and have demonstrated that no two AAVs are mutually immune orthogonal. However, with a known HLA genotype, it may be possible to define a personalized regimen of immune orthogonal AAVs using currently defined serotypes. For instance, use of AAV5 minimizes immune cross-reactivity in mice and non-human primates, as demonstrated by a recent study in which chimeric-AAV5 immunized mice and non-human primates successfully received a second dose of treatment with AAV1^59^.

However, in the human setting we predict that there may be substantially more immune overlap between AAV5 and other AAVs. Additionally, it has been shown that memory B-cells heavily contribute to the antibody response to similar but not identical antigens^65^, indicating that partial orthogonality may not be sufficient. Our analysis suggests that creating a pair of globally orthogonal AAV capsids for human application would require >10 mutations in one of the two proteins. This hypothetical orthogonal AAV capsid presents a substantial engineering challenge, as it requires mutating many of the most conserved regions to achieve immune orthogonality.

Previous work has identified that MHC affinity is highly dependent on anchor residues at either end of the binding pocket^66^. Residue diversity is more tolerated in the center of the binding pocket, though it may be these residues that most impact antigen specificity, as it is thought that they are central to interaction with the T-cell receptor (TCR). Comparing the number of orthologous pairs in 9-mer space with the number of predicted orthologous pairs based on class II binding predictions suggests that only approximately 65% of 9-mer peptides serve as appropriate MHC class II binding cores, even across the thousands of HLA-2 combinations we explore here. This under-sampling of peptide space by MHC molecules likely reflects the requirement for hydrophobic anchor residues and leaves some space for protein de-immunization by mutation of immunogenic peptides to ones which never serve as MHC binding cores. Achieving this while preserving protein function however, has proven difficult even for few HLA alleles, and remains a major protein engineering challenge. New technologies for directly measuring TCR affinity with MHC-presented antigens^67^ will also further clarify the key antigenic peptides contributing to the immune response, and will be useful to inform approaches here.

We also note some limitations to our work. Mainly, we have used inbred C57BL/6J as our mice model, which have very limited MHC diversity^68^, and might not recapitulate other human immunological features, such as differences in antigen processing and presentation. In this regard, we attempted to measure the T-cell response with the IFN-γ ELISPOT assay for a subset of predicted MHCI and MHC II peptides. Concordant with a recent study of pre-existing immunity to Sp-and SaCas9 in humans^38^, we observed a CD4+ T-cell response against some MHCII peptides with mice injected with SaCas9, but not SpCas9. Although we note that the C57BL/6J mice did not show robust responses in general to the AAV-CRISPRs (**Figure S4**), the detection of antibody and T-cell responses to both Cas9^38^ (only SaCas9 in the case of T-cells) and AAVs^1,52^ in humans demonstrates that the immune consequences of treatment will be critical for success in human therapy. Moving forward, this work can be potentially repeated using other mouse models, such as mice expressing human HLA allotypes, however, these models come with their own technical challenges, such as restricted HLA alleles (representing only main MHC II subgroups) as well as a restricted TCR repertoire^68^. In addition, B-cell epitopes can also be predicted and incorporated into immune orthogonality analysis. However, since B-cell epitopes may be both linear and conformational, these are more difficult to predict. Advances and further validation of these *in silico* models will allow for better predictions in the future^69–73^. Finally, recent work has indicated that MHC class I peptides may have significant contribution from spliced host and pathogen-derived peptides created by proteasomal processing^74^. It is unclear how this may affect cross-recognition of proteins we predict to be immune orthogonal. On the one hand, it provides a mechanism whereby very short antigenic sequences spliced to the same host protein may result in cross-recognition of substantially different foreign antigens, however, we expect this to be unlikely due to the massive number of possible spliced peptides between the antigen and entire host proteome.

Overall, we believe our framework provides a potential solution for efficacious gene therapy, not solely for Cas9-mediated genome engineering, but also for other protein therapeutics that might necessitate repetitive treatments. Although using this approach still requires mitigating the primary immune response, particularly CTL clearance, we expect that epitope deletion and low-immunogenicity delivery vectors such as AAVs will mitigate this problem, and the potential for repeated dosage will reduce the need for very high first-dose efficiency.

## ACKNOWLEDGEMENTS

We thank members of the Mali lab for advice and help with experiments, and the Salk GT3 viral core for help with AAV production. This work was supported by UCSD Institutional Funds, the Burroughs Wellcome Fund (1013926), the March of Dimes Foundation (5-FY15–450), the Kimmel Foundation (SKF-16–150), and NIH grants (R01HG009285, RO1CA222826). A.M. acknowledges a graduate fellowship from CONACYT and UCMEXUS. A.M., N.P., and P.M. have filed patents based on this work.

## METHODS

### Computational Methods

For Cas9, we chose 91 orthologs cited in exploratory studies cataloguing the diversity of the Cas9 protein^75^, including several that are experimentally well-characterized. For AAVs, we analyzed 167 sequences, focusing in on all 13 characterized human serotypes, as well as one isolate from rhesus macaque (rh32), one engineered variant (DJ), and one reconstructed ancestral protein (Anc80L65). We then compared total sequence similarity (immunologically uninformed) as well as predicted binding to class I and class II MHC molecules (immunologically informed) between these proteins. Immunologically uninformed sequence comparison was carried out by checking a sliding window of all contiguous k-mers in a protein for their presence in another protein sequence with either zero or one mismatch. Immunologically informed comparison was done in a similar fashion, but using only those k-mers predicted to bind to at least one of 81 HLA-1 alleles using netMHC 4.0^76^ for class I (alleles can be found at http://www.cbs.dtu.dk/services/NetMHC/MHC_allele_names.txt), and at least one of 5,620 possible MHC II molecules based on 936 HLA-2 alleles using netMHCIIpan 3.1^77^ for class II (alleles can be found at http://www.cbs.dtu.dk/services/NetMHCIIpan-3.1/alleles_name.list). We compared the use of netMHC to alternative immune epitope prediction platforms such as the Immune Epitope Database (iedb.org)^78^ and found very strong agreement across software. Ultimately, we chose netMHC because of the larger number of HLA alleles it supports. Sequences were defined as binding if the predicted affinity ranked in the top 2% of a test library of 400,000 random peptides as suggested in the software guidelines. Generation of immune orthogonal cliques was carried out using the Bron-Kerbosch algorithm. Briefly, a graph was constructed with each ortholog as a vertex, where the edges are defined by the number of shared immunogenic peptides between the connecting vertices. Sets of proteins for which every pair in the set is immune orthogonal constitutes a clique. Phylogenetic distance between protein sequences was measured using the BLOSUM 62 matrix excluding indels.

### Experimental Methods

#### AAV Production

AAV2/8, AAV2/2, AAV2/5, AAV2/DJ virus particles were produced using HEK293T cells via the triple transfection method and purified via an iodixanol gradient^79^. Confluency at transfection was between 80% and 90%. Media was replaced with pre-warmed media 2 hours before transfection. Each virus was produced in 5 × 15 cm plates, where each plate was transfected with 7.5 μg of pXR-capsid (pXR-8, pXR-2, pXR-5, pXR-DJ), 7.5 of μg recombinant transfer vector, and 22.5 μg of pAd5 helper vector using PEI (1ug/uL linear PEI in 1× DPBS pH 4.5, using HCl) at a PEI:DNA mass ratio of 4:1. The mixture was incubated for 10 minutes at RT and then applied dropwise onto the media. The virus was harvested after 72 hours and purified using an iodixanol density gradient ultracentrifugation method. The virus was then dialyzed with 1× PBS (pH 7.2) supplemented with 50 mM NaCl and 0.0001% of Pluronic F68 (Thermo Fisher) using 100kDA filters (Millipore), to a final volume of ∼1 mL and quantified by qPCR using primers specific to the ITR region, against a standard (ATCC VR-1616).

*AAV-ITR-F: 5’-CGGCCTCAGTGAGCGA-3’ and*

*AAV-ITR-R: 5’-GGAACCCCTAGTGATGGAGTT-3’*.

#### Animal studies

All animal procedures were performed in accordance with protocols approved by the Institutional Animal Care and Use Committee (IACUC) of the University of California, San Diego. All mice were acquired from Jackson labs. AAV injections were done in adult C57BL/6J mice (10 weeks) through retro-orbital injections using 1×10^12^ vg/mouse.

##### ELISA

*PCSK9:* Levels of serum PCSK9 were measured using the Mouse Proprotein Convertase 9/PCSK9 Quantikine ELISA kit (R&D Systems) according to manufacturer’s guidelines. Briefly, serum samples were diluted 1:200 in Calibrator diluent and allowed to bind for 2 h onto microplate wells that were precoated with the capture antibody. Samples were then sequentially incubated with PCSK9 conjugate followed by the PCSK9 substrate solution with extensive intermittent washes between each step. The amount of PCSK9 in serum was estimated colorimetrically using a standard microplate reader (BioRad iMark).

*Cas9 and AAV:* Recombinant SpCas9 protein (PNA Bio, cat. no. CP01), or SaCas9 protein (ABM good, cat no. K144), was diluted in 1× coating buffer (Bethyl), and 0.5 μg was used to coat each well of 96-well Nunc MaxiSorp Plates (ab210903) overnight at 4 °C. For AAV experiments, 10^9^ vg of AAV-2, -5, -8 or-DJ in 1× coating buffer was used to coat each well of 96-well Nuc MaxiSorp Plates. Plates were washed three times for 5 min with 350 μl of 1× Wash Buffer (Bethyl) and blocked with 300 μl of 1× BSA Blocking Solution (Bethyl) for 2 h at RT. The wash procedure was repeated. Serum samples were added at 1:40 dilution, and plates were incubated for 5 h at 4 °C with shaking. Wells were washed three times for 5 min, and 100 μl of HRP-labeled goat anti-mouse IgG1 (Bethyl; diluted 1:100,000 in 1% BSA Blocking Solution) was added to each well. After incubating for 1hr at RT, wells were washed four times for 5 min, and 100 μl of TMB Substrate (Bethyl) was added to each well. Optical density (OD) at 450 nm was measured using a plate reader (BioRad iMark).

#### Epitope prediction and peptide synthesis

The MHC-binding peptides for our mouse model were predicted using the netMHC-4.0 and netMHCIIpan-3.1 online software with the alleles H-2-Db and H-2-Kb for class I and H-2-IAb for class II. For MHCII, the top 10 peptides for Sp-and SaCas9 and top 5 peptides for AAV-8 and AAV-DJ by percentile binding were selected for synthesis by Synthetic Biomolecules as crude materials. For MHCI, we selected the top 20 peptides for Sp-and SaCas9 and the top 10 for AAV-8 and AAV-DJ. All peptides were dissolved in DMSO with a concentration of 40 mg ml^−1^ and stored at −20 °C.

#### IFN-γ ELISPOT assay

CD8+ T cells were isolated from splenocytes using magnetic bead positive selection (Miltenyi Biotec) 6 weeks after virus infection. A total of 2 × 10^5^ CD8+ T cells were stimulated with 1 × 10^5^ LPS-blasts loaded with 10 μg of individual peptide in 96-well flat-bottom plates (Immobilon-P, Millipore) that were coated with anti-IFN-γ mAb (clone AN18, Mabtech) in triplicate. Concanavalin A (ConA) was used as positive control. After 20 h of incubation, biotinylated anti-mouse IFN-γ mAb (R4–6A2; Mabtech), followed by ABC peroxidase (Vector Laboratories) and then 3-amino-9-ethylcarbazole (Sigma-Aldrich) were added into the wells. Responses are expressed as number of IFN-γ SFCs per 1 × 10^6^ CD8+ T cells.

**Supplementary Figure 1.**
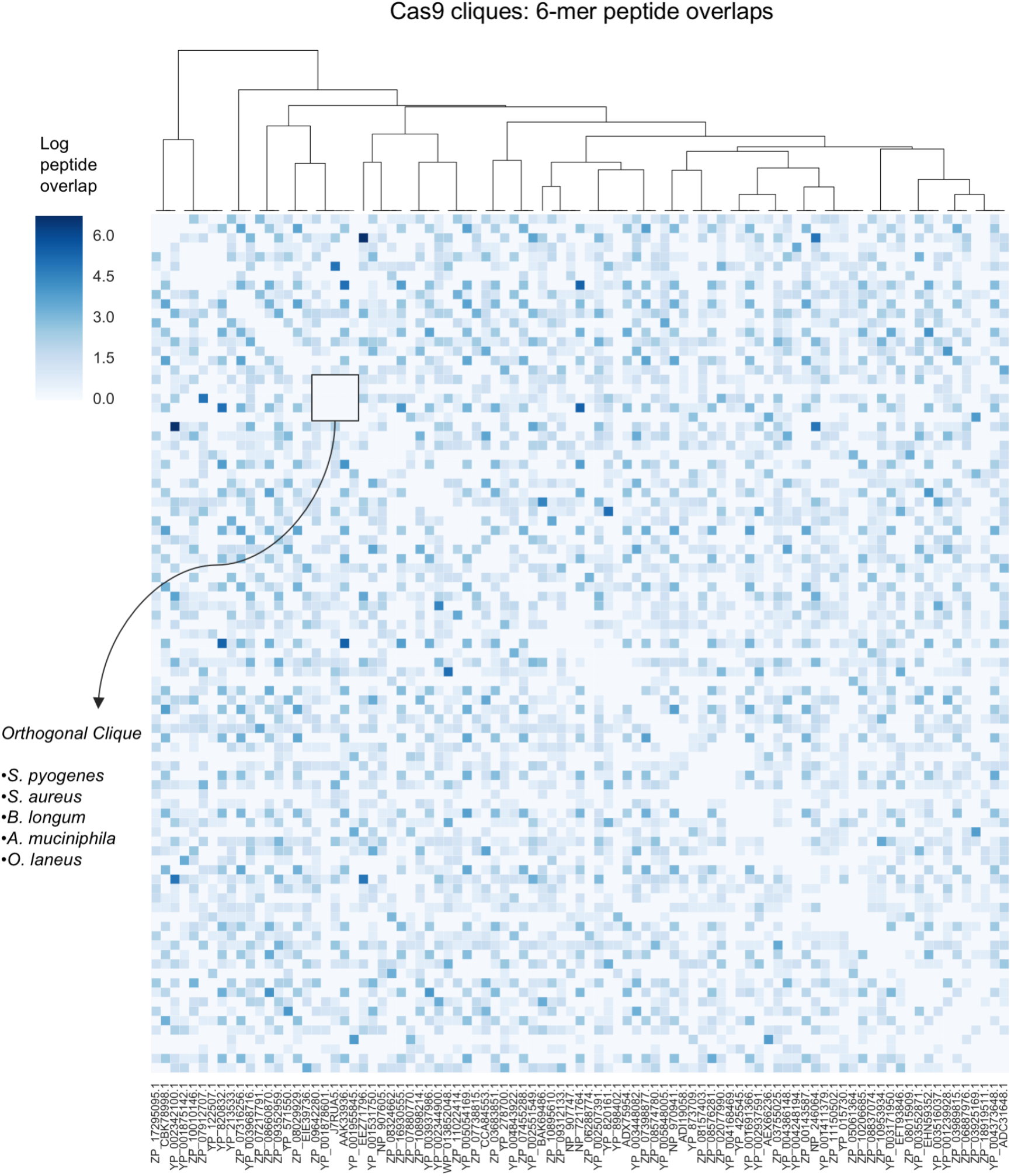
Cas9 immune orthogonal cliques. Cliques corresponding to 6-mer overlaps are depicted. An example of an orthogonal clique is highlighted, which includes Cas9s from: *S. pyogenes, S. aureus, B. longum, A. muciniphila*, and *O. laneus*.

**Supplementary Figure 2.**
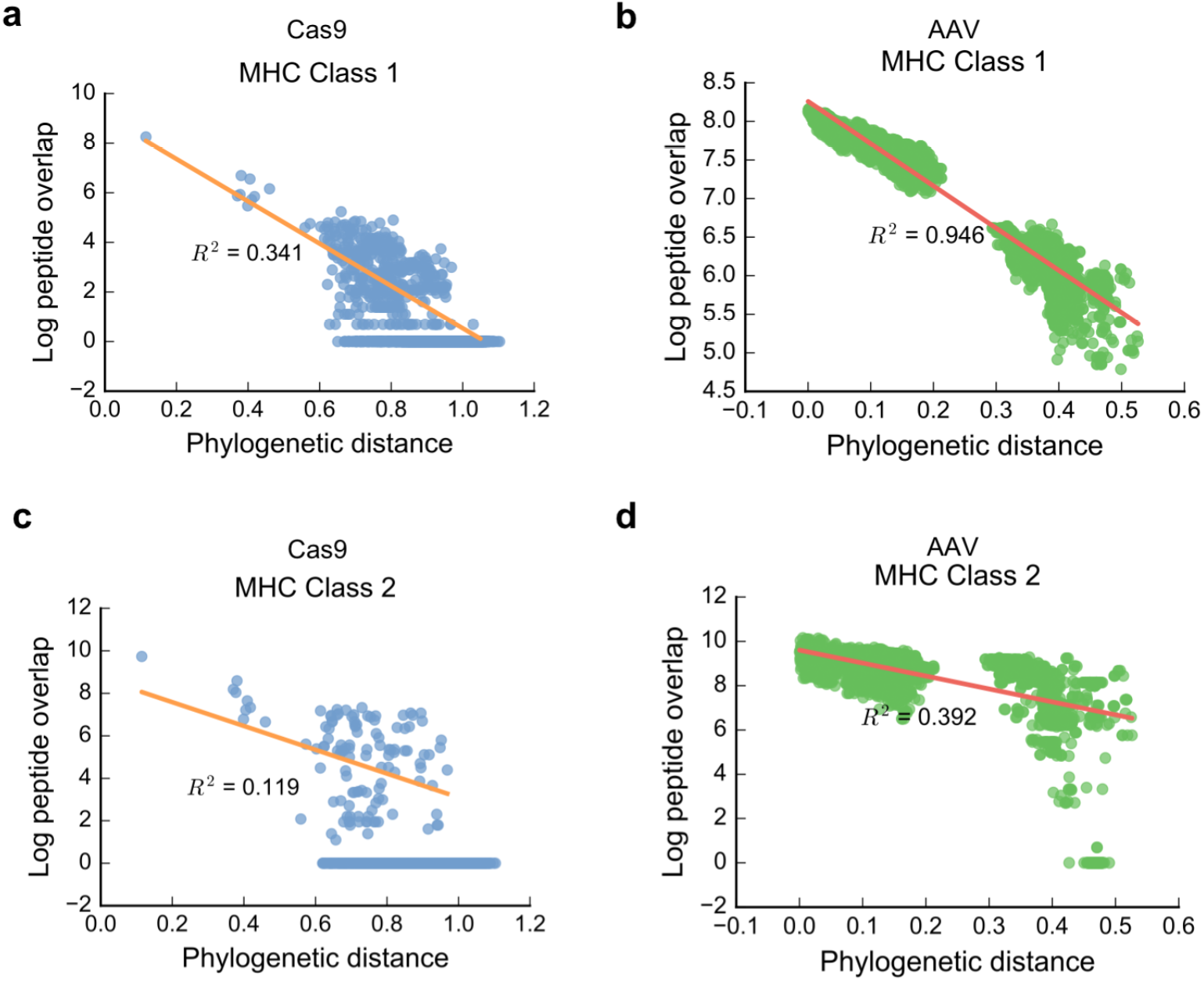
In silico analyses and comparisons of immunogenicity of Cas9 and AAV orthologs. Linear regressions exclude pairs with no overlap. **(a)** Cas9 MHC class I peptide overlap vs. phylogenetic distance. **(b)** AAV MHC class I peptide overlap vs. phylogenetic distance. **(c)** Cas9 MHC class II peptide overlap vs. phylogenetic distance. **(d)** AAV MHC class II peptide overlap vs. phylogenetic distance.

**Supplementary Figure 3.**
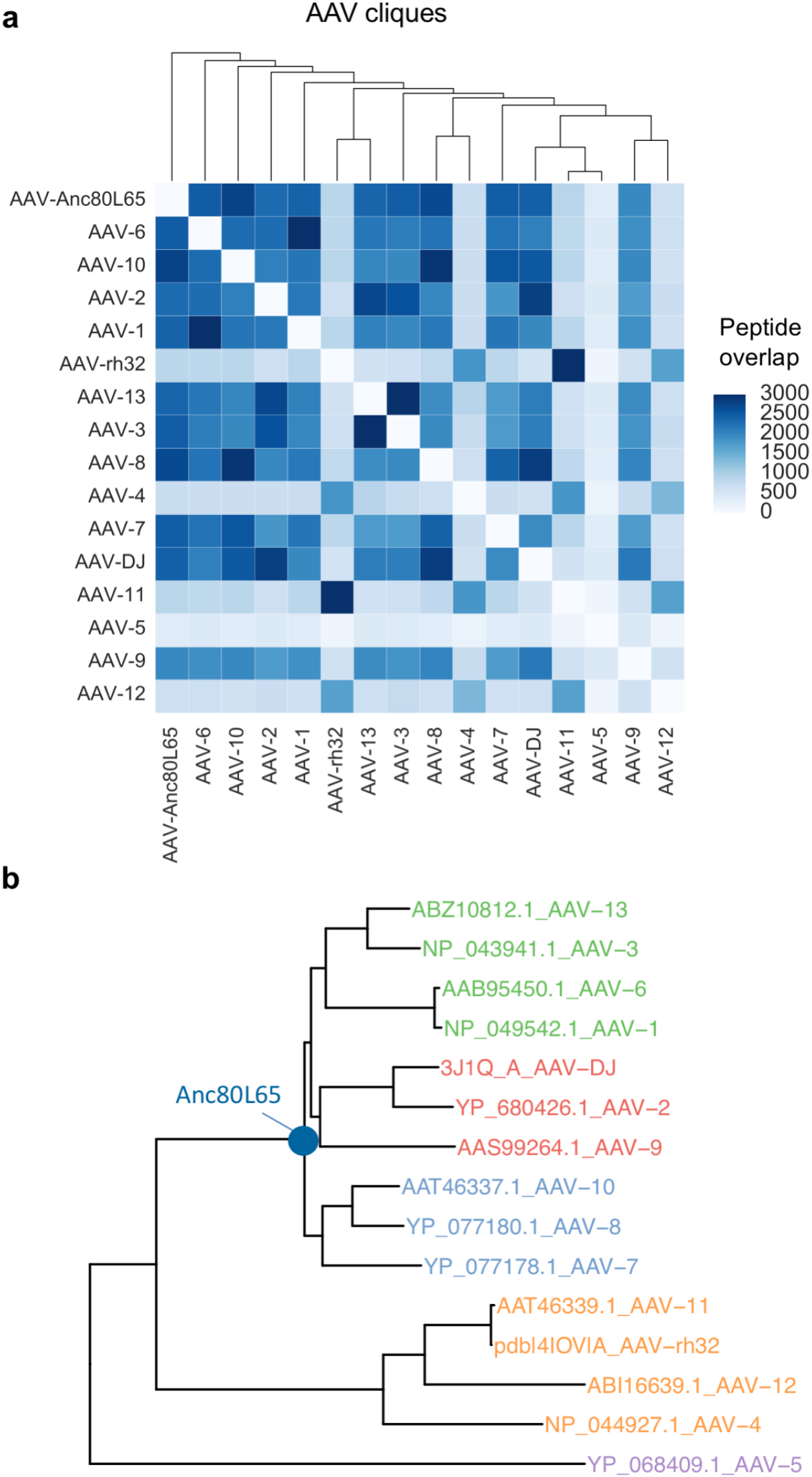
Major AAV serotype groups. **(a)** AAV immune orthogonal cliques over 81 HLA alleles. AAV5 is the most immune-divergent in comparison to the other serotypes. No orthogonal cliques exist. **(b)** AAV phylogeny showing major serotype groupings as well as the position of the reconstructed sequence Anc80L65.

**Supplementary Figure 4.**
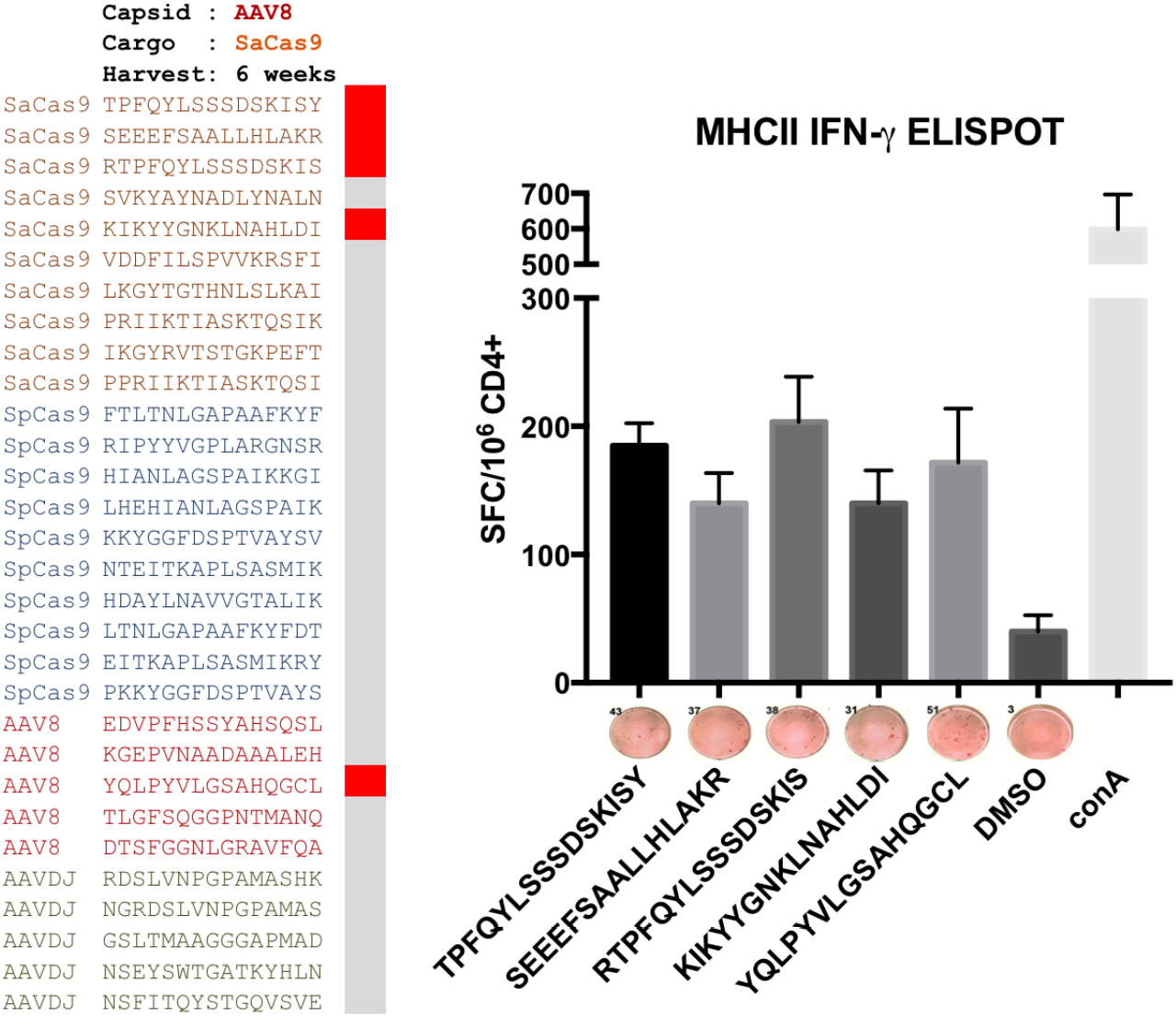
Experimental validation of a MHCII peptide predictions via IFN-γ ELISPOT. Mice were injected retro-orbitally with 10^12^ vg/mouse of AAV8-SaCas9 targeting the PCSK9 gene and were sacrificed after 6 weeks (n=6). Purified CD4+ T cells from splenocytes were seeded at 2×10^5^ cells per well in triplicate. 1×10^5^ lipopolysaccharide-activated antigen presenting cells (APCs) from control mice were added to each well. Cells were incubated with highly immunogenic MHC-II predicted peptides for 20h. Spots were developed with biotinylated anti-IFN-γ. A one-way ANOVA with post hoc Dunnett’s test was performed to determine statistical differences with DMSO for all peptides (left panel), and the significant peptides were plotted (right panel).

